# C19ORF66 is an Interferon-Stimulated Gene (ISG) which Inhibits Human Immunodeficiency Virus-1

**DOI:** 10.1101/050310

**Authors:** Weidong Xiong, Deisy Contreras, Joseph Ignatius Irudayam, Ayub Ali, Otto O. Yang, Vaithilingaraja Arumugaswami

## Abstract

Innate immunity is the first line of defense against invading microbes^1^. The type I interferon (IFN) pathway plays a key role in controlling Human Immunodeficiency Virus type 1 (HIV-1) replication^2,3^. We identified an IFN-α stimulated gene C19ORF66 that we term Suppressor of Viral Activity (SVA). Full length SVA-1 protein inhibits HIV-1 by blocking virion production. SVA splice variants truncated at the C-terminus and/or disrupted at the nuclear export signal (NES) lose antiviral activity and localize to nucleus, while full length SVA-1 co-localizes with HIV-1 p24 protein in the cytoplasmic compartment of infected cells. SVA-1 is structurally and functionally conserved across species, including mouse and chimpanzee. We provide the first description of the effector function of the gene SVA/C190RF66 as an innate immune factor with anti-HIV-1 activity.

## INTRODUCTION

Innate immune responses are fast acting, first lines of defense against various forms of microbial infection, comprised of pathogen recognizing receptors (PRR) and effectors for generating inflammatory and antimicrobial responses. Detection of viral pathogen associated molecular motifs (PAMP) by PRR such as toll-like receptor-3 (TLR-3), RIG-I, MDA5, DAI, cGAS and STING prompts the activation of downstream factors including NF-κB, c-Jun/ATF-2, CBP/P300, interferon regulatory factor 3 (IRF3) and IRF7 resulting in expression of inflammatory, and type I interferon (IFN) α and β cytokines. These IFNs subsequently induce an antiviral state through autocrine and paracrine mechanisms by engaging type I IFN receptors and activating the JAK/STAT signaling cascade initiating the formation of the IFN-stimulated gene factor 3 (ISGF3) complex. In the nucleus, ISGF3 complex binds to IFN-stimulated response elements (ISRE) promoting the transcription of hundreds of interferon-stimulated genes (ISGs) that are involved in antiviral effector function^10^.

Type I IFN signaling plays important role in the pathogenesis of Human Immunodeficiency Virus type 1 (HIV-1) infection^2,3,11^. Reverse transcribed proviral DNA is recognized by cGAS to trigger production of type I IFNs^12^. Virus-dependent upregulation of ISGs including IS15, MX1, OAS3, and IRF7 occurs in infected CD4^+^ T cells^11^. Several cellular antiviral effector ISGs can interfere with various stages of HIV-1 replication, including CNP, TRIM5α, APOBEC3G, tetherin (CD317), SAMHD1, MX2 and IFI16 ^13^^-^^19^.
HIV-1 evolved strategies to evade many of these factors through its proteins such as Vif, Vpu and capsid ^13,20,21^.

Here, we identify a new ISG C19ORF66 (synonym: FLJ11286) which codes for an uncharacterized protein UPF0515 of previously unknown function. We describe the novel effector function of C19ORF66 as an inhibitor of HIV-1 replication.

## METHODS

### Cells

The human embryonal kidney (HEK)-293-FT cell line (Life Technologies, CA, USA), and HeLa cell line (American Type Culture Collection, VA, USA) were maintained on complete Dulbecco’s modified Eagle’s medium (DMEM) (Sigma Aldrich, MO, USA) in a humidified cell culture incubator with 5% CO^2^ at 37°C. Complete DMEM comprised of 10% fetal bovine serum (FBS), 10 mM Hepes, 10 mM nonessential amino acids, penicillin (100 units/ml), streptomycin (100 mg/ml), and 2 mM L-glutamine (Life Technologies, USA).

### IFN-α treatment

HeLa cells and de-identified human primary cells, peripheral blood mononuclear cells (PBMC) and macrophages (CFAR Virology Core Laboratory at UCLA AIDS institute, CA, USA), were used. Cells (2x10^5^ per well) were plated into 12-well plate and next day morning stimulated with human IFN-α (R and D Systems, MN, USA) at a concentration of 5000 IU/ml for indicated time points. Untreated cells were included as negative control. Cells were collected for gene expression studies by Western blotting and reverse transcription-quantitative PCR (RT-qPCR).

### Plasmids

A self-inactivating (SIN), third generation lenti vector plasmid was used. The individual gene inserts were sub-cloned into lenti vector using with *Age*I-*Xho*I or *Age*I-*Pac*I restriction digestions at upstream of internal ribosome entry site (IRES) sequence and following selection marker mCherry or puromycin resistance gene (*pac*). The sequences of the primers used for gene amplification is provided in Extended Data Table 1. For introducing mutations or deletion, a two-step polymerase chain reaction (PCR)-based approach was used. The complementary DNA (cDNA) of each gene was either procured (Open Biosystems, GE Dharmacon, CO, USA) or amplified by revers-transcription PCR from HeLa cells or H9 human embryonic pluripotent stem cell-derived hepatic lineage cells^10^. The cDNA was generated for full length C19ORF66 (NCBI accession ID: NM_018381.3), a novel C19ORF66 splice variant 5 (isoform X4) and an additional C19ORF66 splice variant XN (sequence ID XM_006722787.1 obtained from NC_000019.10 Chromosome 19 Reference GRCh 38 Primary Assembly in 2014). C19ORF66 nuclear localization signal (133-<u>RK</u>C<u>RKR</u>-138) was abolished by introducing point mutations (<u>AA</u>C<u>AAA</u>) by two-step mutagenesis PCR. The NES mutant was generated by deleting nucleotides coding for amino acids 261-LEDLDNLILED-271. The control lentivector contained luciferase gene or multiple cloning site (MCS). The expression of bicistronic gene cassette was driven by elongation factor-1 alpha (EF-1 alpha) cellular promoter. For packaging lenti vectors, plasmids psPAX2 [a gift from Didier Trono (Addgene plasmid # 12260)] and pCMV-VSV-G [a gift from Bob Weinberg (Addgene plasmid # 8454)] were used^30^. The psPAX2 plasmid codes for HIV proteins (Gag, Pol, Tat and Rev) and cis-elements (RRE and PPTc). Vesicular Stomatitis Virus-glycoprotein (VSV-G) envelop gene is coded by pCMV-VSV-G plasmid, which is used for production of VSV-G pseudotyped lenti viral particles. A plasmid containing HIV-1 NL4-3 proviral genomic DNA was for wild-type HIV-1 experiments.

### RT-qPCR Analysis

Total RNA was extracted from cells by using the RNeasy RNA extraction kit (QIAGEN, USA) as per manufacturer’s instruction. RNA samples were reverse-transcribed using SuperScript III Reverse Transcriptase kit (Life Technologies, USA) with random primers as described by the manufacturer. Quantification of gene expression was done using Platinum SYBR Green qPCR SuperMix-UDG with ROX Kit (Life Technologies) by the ViiA7 real-time PCR system (Applied Biosystems). Housekeeping human gene GAPDH was used as an internal reference control. The sequences of qPCR primer are given in Extended Data Table 2. Quantitative PCR was done under the following conditions: 50°C for 2 min; 95°C for 2 min followed by 40 cycles of 95°C for 15 sec and 60°C for 1 min. To provide comparison among various samples, the Ct value of each gene is normalized to that of the housekeeping gene GAPDH.

### Lentiviral Vector Packaging and viral particle production

Lentiviral vector packaging was done using HEK-293-FT (Life Technologies, USA) cells as follows. 3x10^6^ cells were plated in a 10-cm dish and 24 hours later, the cells were transfected with lentiviral vector (3.2 μg), psPAX2 (3 μg) and pCMV-VSV-G (1 μg) plasmid DNA using polyethyleneimine (PEI, 25 kDa) (Polysciences, Inc., PA, USA) transfection solution (DNA to PEI ratio of 1:5). At 72 hours post-transfection, cell-free culture supernatant was collected for measuring viral particle production (HIV p24 quantitation) and subsequent round of transduction. At the same time, the cells were also harvested for flow cytometry analysis and intra-cellular p24 protein measurement. Cells were lysed in 1x Molecular cell lysis buffer (Cell Signaling Technology Inc., MA, USA) comprised of 1 mM phenylmethanesulfonyl fluoride (Sigma Aldrich, MO, USA), protease inhibitor cocktail (Roche Diagnostics Corporation, IN, USA) and 0.5% triton X-100. HIV p24 (ng/ml) measurement was performed in CFAR Virology Core Laboratory at UCLA AIDS institute using HIV-1 p24 ELISA kit (PerkinElmer Life Sciences, Inc., MA, USA) as per manufacturer’s protocol.

### Lentiviral Vector Transduction and Flow Cytometry Analysis

The HEK-293FT cells (2x10^4^ cells/well) were seeded into 48-well plate and at 24 hours post-plating, 0.5 ml of cell-free culture supernatant containing lentiviral particles were added to each well. 24 hours post-transduction, media were replaced with fresh complete DMEM; and at 72 hours post-transduction, the cells were collected and fixed with buffered formalin (2.5%). BD Fortessa flow cytometer (BD Biosciences) was used for data acquisition (mCherry expressing cells) using FACSdiva software. Data were analyzed using FlowJo software (TreeStar Inc., USA).

### Human Immunodeficiency Virus-1 replication experiment

HEK-293T cells were used for transfection. Briefly, 0.8x10^6^ cells were seeded in a well of 6 well-plate at one day before transfection to reach 60 - 75% confluency in the day of transfection. The HIV-1 plasmid pNL4-3 (2 μg per well) with vector control plasmid or C19ORF66 expressing plasmid (0.5 μg per well) were transfected in triplicates with BioT kit (Bioland Scientific, Paramount, CA). At 48 hour post-transfection, the cells and cell-free supernatants were collected for HIV-p24 ELISA assay.

### Immunocytochemistry and confocal Imaging

HeLa cells (2x10^4^ per well) were seeded into Lab-Tek II Chamber Slide (Thermo scientific, USA). The next day, the cells were transfected with each plasmid construct (200 ng per well) using Lipofectamine 2000 (Life Technologies, USA). At 3 day post transfection, the cells were fixed in 4% paraformaldehyde for 20 mins at 4°C. Following five time of phosphate buffered saline (PBS) washes, the cells were blocked using dilution buffer (10% fetal bovine serum, 3% BSA, 0.1% Triton-x 100 in PBS) and incubated with primary antibody (1:200 dilution) for 5 hours to overnight at 4°C. Subsequently, the cells were washed five times with PBS. The secondary antibody (1:1,000 dilution) was added and incubated for 1 hour at room temperature. Subsequently, the cells were subjected to five time PBS washes. Primary antibodies, including mouse monoclonal anti-HIV1 p24 antibody [39/5.4A] (Abcam, USA), rabbit polyclonal anti-HA antibody and mouse monoclonal anti-flag antibody (Clone M2) (Sigma Aldrich, USA) were used. Secondary antibodies - goat anti-mouse IgG conjugated to Alexa Fluor 488 or 594, goat anti-rabbit IgG conjugated to Alexa Fluor 488, and Donkey anti-Rabbit IgG conjugated to Alexa Fluor 647 were used (Life Technologies, USA). Immunostained slides were mounted with coverslips using ProLong Gold antifade reagent containing DAPI (Life Technologies, USA) and examined by fluorescence microscopy. Confocal images were collected on a Nikon A1R inverted laser scanning confocal microscope and analyzed with ImageJ software.

### Western Blotting

Cells were directly lysed with 1x Molecular cell lysis buffer (non-denaturing), or Laemmli sample buffer (BIO-RAD, CA, USA) containing 5% β-mercaptoethanol (Sigma Aldrich, USA). The cell lysates with Laemmli sample buffer were boiled for 5 minutes in a heat block. Protein lysates (15-20 μg/lane) were separated using 4-15% gradient sodium dodecyl sulfate-polyacrylamide gel (BIO-RAD, USA) and transferred onto nitrocellulose membranes (BIO-RAD, USA). Membranes were blocked using 5% nonfat milk and 0.05% of Tween-20 in PBS (blocking buffer) and then incubated with appropriate primary antibody for overnight at 4°C. Primary antibodies, mouse monoclonal anti-HIV1 p24 antibody [39/5.4A], rabbit polyclonal anti-HA antibody (Sigma Aldrich, USA), rabbit polyclonal anti-C19orf66 antibody (Sigma Aldrich, USA), and mouse monoclonal anti-β-actin antibody (Sigma Aldrich, USA), were diluted in blocking buffer (1 in 1,000 dilution). The blots were washed three times with 0.1% of Tween-20 containing PBS and subsequently incubated with the goat secondary antibody (anti-mouse or anti-rabbit) conjugated to horseradish peroxidase (HRP) (1:5000 dilution) for 1 hour at room temperature with shaking. The membranes were incubated with SuperSignal West Pico chemiluminescent solution (Thermo Scientific, USA) for 5 minutes at room temperature. ChemiDoc XRS+ System with Image Lab™ Software (BIO-RAD, USA) was used for blot image acquisition and analysis.

### Statistical analysis

Error bars in the graph depict the standard deviation. P-values were calculated by the two-tailed student’s t-test and significance was indicated if p value is p<0.05 (*); p<0.001 (**); p<0.0001 (***).

## RESULTS

### C19ORF66 is an ISG that inhibits HIV-1 production

Unbiased RNA sequencing analysis of IFN-α stimulated cells revealed the induction of C19ORF66. This gene and its promoter showed cis-element binding sites for transcription factors IRF1, IRF7A and STAT1, confirming it as a downstream target of IFN signaling. Moreover this locus in chromosome 19 is flanked by genes involved in inflammatory and IFN signaling pathways such as TYK2, ANGPTL6, ICAM1, ICAM3 and ICAM4, further suggesting an important role in innate and inflammatory responses.

Whole PBMC and blood monocytes showed significant upregulation of C19ORF66 mRNA following 6 hours of IFN-α treatment (Fig. 1a), and HeLa cells showed corresponding expression of endogenous 35 kDa C19ORF66 protein at 6 hours as well as a minor isoform species of 30 kDa after 24-48 hours (Fig. 1b). When HIV-1-based lentiviral vectors were constructed to deliver genes for C19ORF66 and other candidate ISGs, viral particle production of the C19ORF66 vector 10-fold reduced compared to the vector control and most other ISG vectors (Fig. 1c). C19ORF66 expression did not exert any cytotoxicity. Flow cytometry analysis of transduced 293-FT cells confirmed the lack of production of lentiviral particles with the C19ORF66 gene (Fig. 1d and Extended Data Fig. 1). This occurrence was also seen for 293T cells transfected with whole wild-type proviral HIV-1 when co-transfected with an expression vector for the C19ORF66 gene, which reduced both intracellular p24 expression and release of p24 virions (Fig. 1e). These findings suggested that C19ORF66, Suppressor of Viral Activity-1, SVA-1, is an antiviral effector in the IFN-α mediated innate immune response.

**Figure 1.**
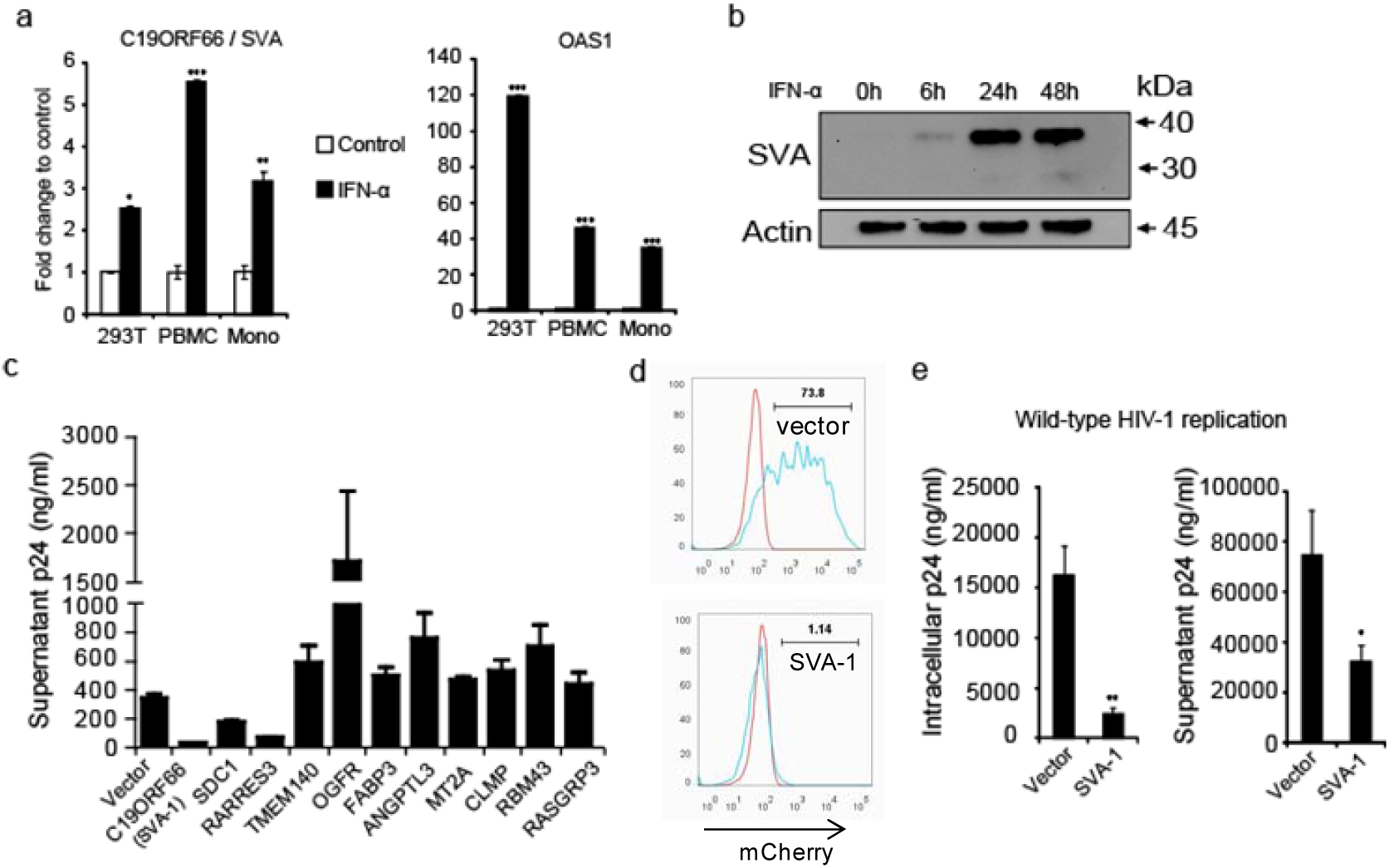
Analysis of expression kinetics and function of IFN stimulated gene C19ORF66 (SVA-1) (a) Graph shows the expression of C19ORF66 transcript in human cells following IFN-α treatment for 6 hours. Known ISG OAS1 expression is tested as control. (b) Expression kinetics of C19ORF66 gene products in HeLa cells at indicated time points post-IFN-α treatment. Western blot analysis shows a prominent 33 kDa protein product and a 28 kDa product suggestive of induction of full length and splice variant of C19ORF66. (d) Screening of candidate ISGs for anti-viral activity. Graph depicts that the C19ORF66 inhibited the production of pseudotyped lentiviral particles as measured by p24 capsid protein released into the 293-FT cell culture supernatant. (d) Flow cytometry analysis of 293-FT cells transduced with control vector-mCherry or C19ORF66-mCherry lentiviruses. Only a small percentage (1.1%) of C19ORF66 transduced cells expressed mCherry indicating lack of functional viral particles generation. Red peak in the histogram indicates the mock transduced negative control cells. (e) Graphs show the wild-type HIV-1 viral production in the presence of SVA-1 in 293-FT cells. HIV-1 p24 level in both intracellular and supernatant was significantly reduced by SVA-1. Representative data from three independent experiments were shown. Mono: monocytes derived from PBMC

### SVA isoforms vary in their antiviral activities and localization patterns

To further characterize SVA-1, we examined its isoforms utilizing pseudotyped lentiviral vector system. Due to alternative splicing event, various protein isoforms were reported in reference database (Extended Data Fig. 2). We identified isoforms X4 (which lacks exon 7 due to a C-terminal truncation) and XN (which contains a longer exon 6 due to alternative splicing to an upstream splice acceptor site) while cloning SVA-1 from IFN-α treated cells. The full length SVA-1 protein contains a nuclear localization signal (RRVPQRKEVSRCRKCRK) at amino acid position 121-137 along with a C-terminal glutamic acid-rich domain (aa 270-291) (Fig. 2a). The first 163 amino acids of the full SVA-1 are shared with the X4 and XN isoforms, but both these variants lack the enriched glutamic acid sequence found at the C-terminal domain (Fig. 2a). Visualization in HeLa cells showed that full length SVA-1 localizes predominantly in the cytoplasm, as opposed to the splice variants that are nuclear localized (Fig. 2a). Further, we tested these variants for antiviral activity and observed that they had decreased efficiency compared to full length SVA-1 (Fig. 2). Full SVA-1 protein strongly co-localized with viral protein p24 in the cytoplasm (Fig. 2e) with occasional cytoplasmic co-localization with Rev and Tat proteins (Extended Data Fig. 3). The data suggest that the SVA-1 C-terminal portion contains a crucial domain that determines not only the localization of the protein but its antiviral activity as well.

**Figure 2.**
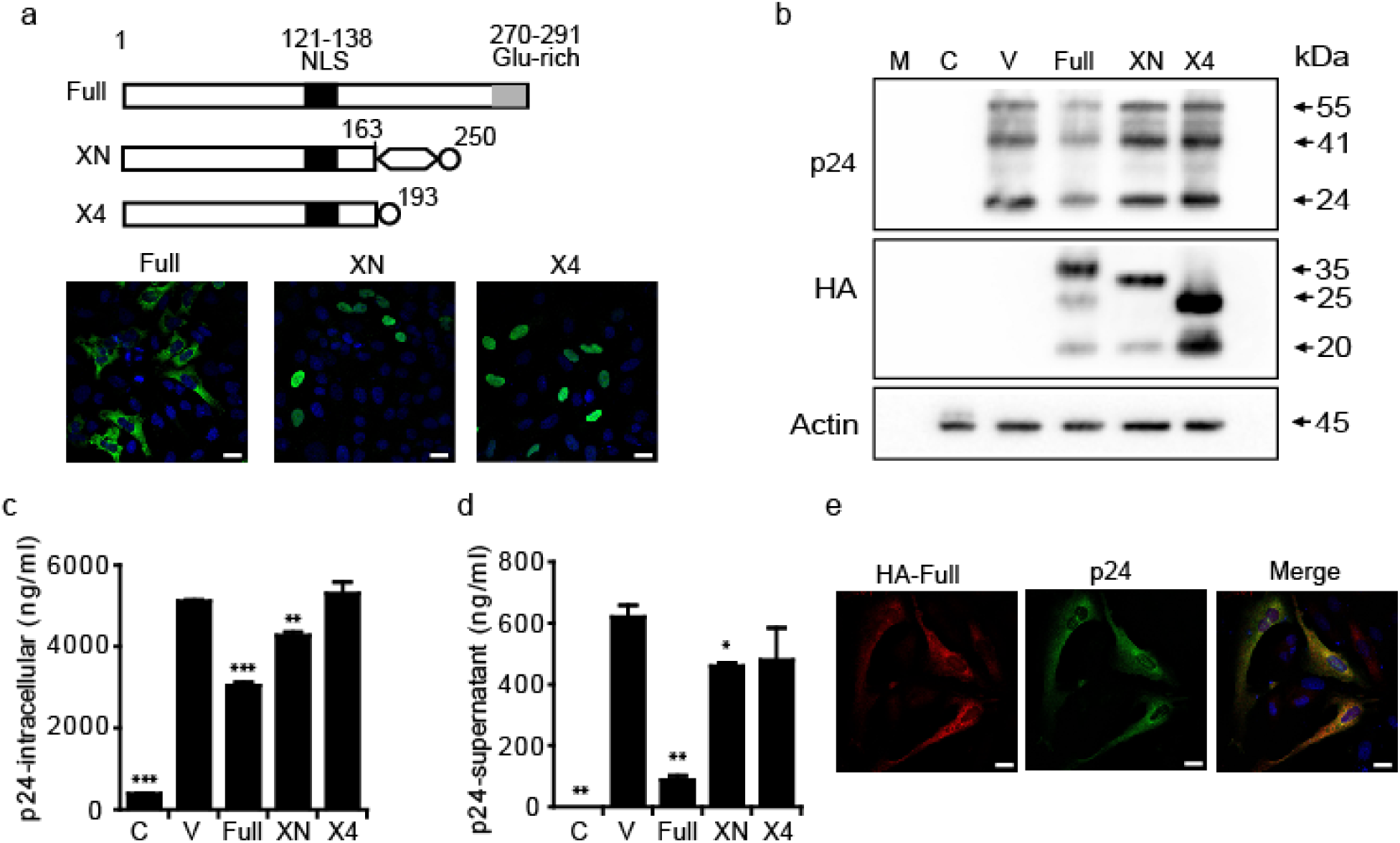
Localization and functional assessment of SVA-1 and its splices variants. (a) Schematic diagram shows predicted protein sequence motifs of full length SVA-1 and splice variants XN and X4. All the indicated isoforms retained the N-terminal 163 amino acids including nuclear localization signal (NLS). Confocal microscopic images show cytoplasmic localization of full length SVA-1 protein and nuclear location of splice variants in transfected HeLa cells. The proteins were immunostained for N-terminal HA epitope. Scale bar: 20 μm. (b, c, d) Packaging assay in 293-FT cells. Lentiviral vectors carrying SVA isoform genes (cDNA) were individually packaged in 293-FT cells. Western blot and ELISA analyzes indicate strong inhibition of HIV p24 protein production by full length SVA-1 in transfected cells. XN isoform has a significant but moderate inhibition of intra-and extra-cellular viral particle (p24 level by ELISA) production. (e) Confocal microscopic analysis indicates the co-localization of full length SVA-1 and HIV p24 proteins in cytoplasm of HeLa cells. Glu-rich: glutamic acid rich domain. Representative data from three independent experiments were shown. M: protein marker; C: non-transfected negative control cells; V: vector control transfected cells.

### Nuclear export is critical for the antiviral function of SVA-1

To assess the mechanism for the reduced antiviral activity of SVA-1 after C terminal truncation, we generated a panel of mutants (Fig. 3). When we assessed the ability of altered gene sequences to block the production of lentiviral vector particles, the data showed that both the full length protein and the altered NLS mutant were able to maintain antiviral function, whereas the mutants without C-terminal amino acids 74 and 104 significantly lost their ability to reduce viral production. Assessing how these alterations affected cellular localization of these proteins, C-terminal truncation also changed the localization to nuclear from cytoplasm, allowing us to identify a critical nuclear export signal (NES sequence 260-<u>L</u>LED<u>L</u>DN<u>L</u>I<u>L</u>E-270) within C-terminal that is required for antiviral activity. Together, these data suggest that the intact C-terminal domain is crucial for the antiviral activity of SVA-1.

**Figure 3.**
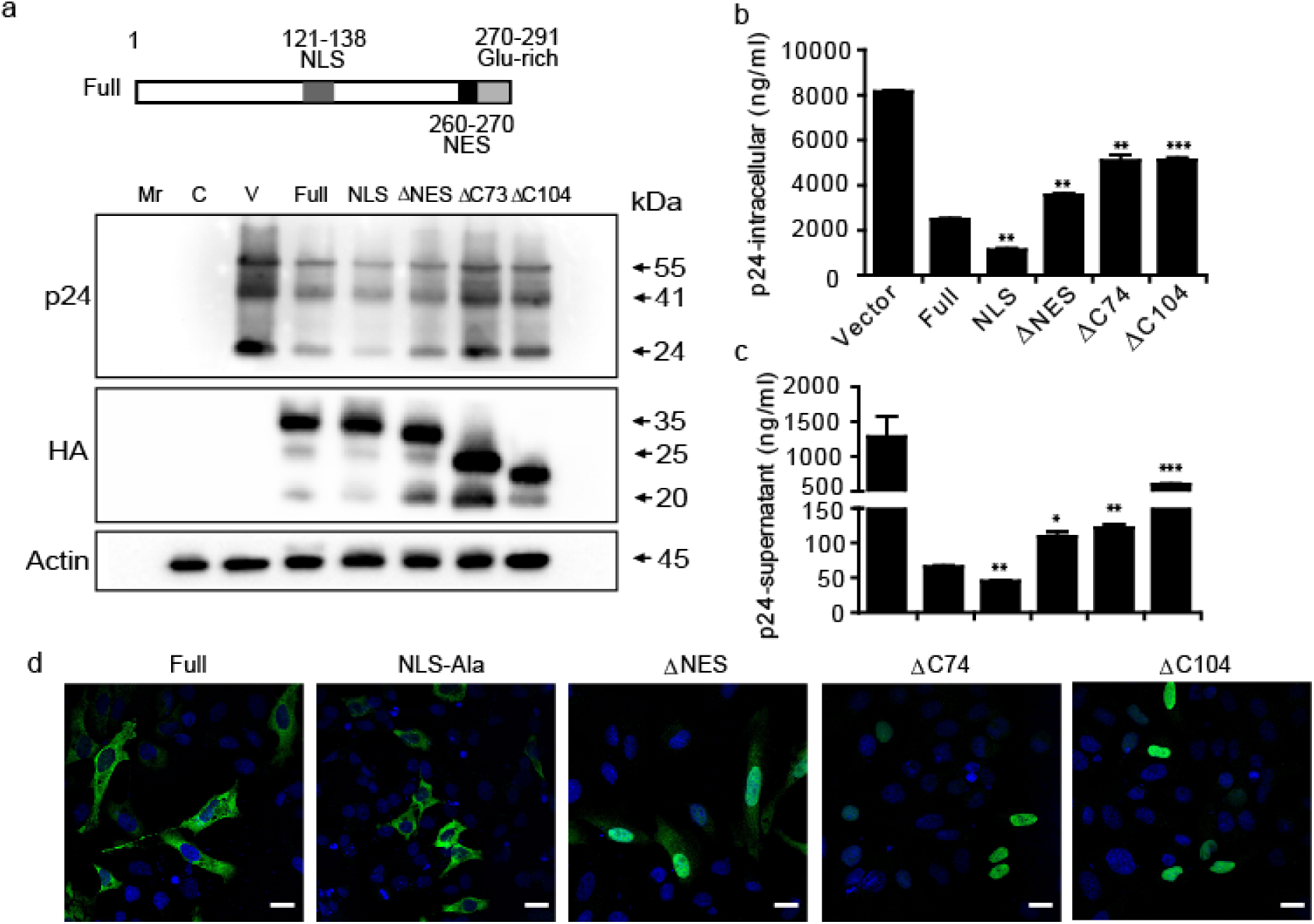
Mapping SVA-1 protein domains essential for antiviral function. Lentiviral vectors expressing SVA-1 mutants lacking NLS, nuclear export signal (NES) and C-terminal 74 or 104 amino acid sequences were packaged individually in 293-FT cells. (a) Western blot analysis shows reduction in HIV p24 protein level in both full length and mutant NLS construct transfected cells. The N-terminal HA-tagged mutant proteins were expressed at the expected protein size. Actin was included as a loading control. (b, c) Graphs show the intra-and extra-cellular levels of HIV p24 protein. The p24 amount, in the transfected cells, was significantly increased with mutants lacking NES and C-terminal domains compared to that of full length. (d) Localization pattern of mutant SVA-1 proteins in HeLa cells. Mutant NLS as well as full length proteins predominantly found in the cytoplasmic compartment. The proteins without nuclear export signal and C-terminal domains localized mostly in the nucleus. Scale bar: 20 μm. Mr: protein marker; C: non-transfected negative control cells; V: vector transfected cells.

### SVA-1 antiviral function is conserved across species

Retroviruses have a broad host range and cells have evolved conserved antiviral strategies to counteract infection. IFN-α is an effective inhibitor of diverse primate and non-primate lentiviruses, including SIV_Mac_, FIV, EIAV, and other retroviruses^22^. The SVA-1 protein is over 95% conserved amongst the mouse, chimpanzee, and human (Extended Data Fig. 4). Given this cross-species conservation, we investigated whether antiviral activity against lentiviral replication was also shared. C19ORF66 (SVA-1) maintained its antiviral property across these species in reducing intracellular viral particle production and release (Fig 4). Taken together, SVA-1 forms a part of a well-conserved innate antiviral response. The SVA-1 protein homologues of murine and primate species can be expected to restrict lenti-or retro-viruses of respective species. Previous studies have shown that ISGs such as cGAS, MX2, and CNP possess broad spectrum anti-retroviral function^12,14,19^.

**Figure 4.**
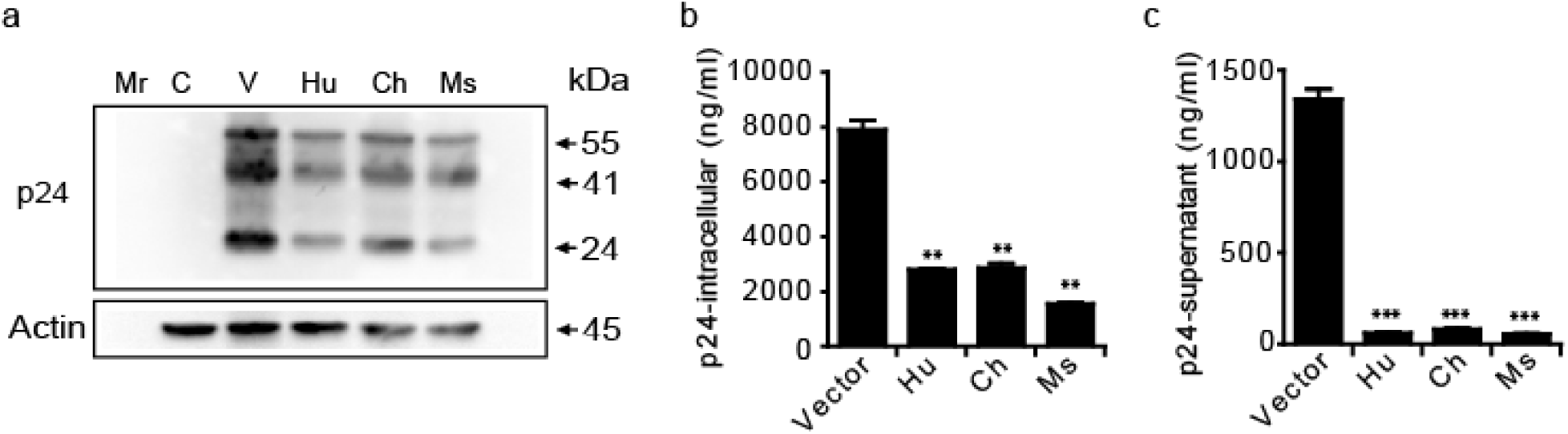
Evaluating the function of SVA-1 homologues. Full length SVA-1 homologues of human, chimpanzee and mouse were tested in a lentiviral packaging assay. The homologues strongly inhibited the lenti viral replication as determined by reduction in p24 protein by western blot (a) and ELISA (b, c).

## DISCUSSION

Inhibition of HIV-1 replication by interferon has been well recognized, however the downstream effector ISGs mediating this activity remain to be fully understood. We utilized both authentic NL-43 wild type HIV-1 and VSV-G pseudotyped HIV-1 based lentivector system to provide a functional annotation of SVA-1. SVA-1 is an interferon-stimulated gene, as its expression was induced in human cells by IFN-α treatment. Previous studies have demonstrated that the IFN-α treatment of macrophages and T cells can block HIV-1 infection various steps ranging from reverse transcription to Env assembly to release of virions^22^^-^^25^. IFN-α treatment predominantly interferes with the accumulation of reverse transcribed HIV-1 cDNA^26,27^. This impairment can be reversed by proteasome inhibition suggesting a key role of the ubiquitin proteasome system ^22^. Some defined mechanisms have included: APOBEC3G, a cytidine deaminase that causes mutations during viral reverse transcription^20^, SAMHD1, which reduces the intracellular pool of deoxynucleotide triphosphate (dNTPs) available for reverse transcription^18^, TRIM5α, which restricts viral capsid uncoating and enhances proteasome-mediated degradation^16^, and CNP, which interferes with virion assembly^14^. Here we add SVA-1 as an additional antiviral ISG.

We confirm SVA-1 to be an interferon-stimulated gene that suppresses HIV-1 replication via impairment of intracellular Gag protein synthesis resulting in a reduced viral particle release. We observe that the cytoplasmic localization of SVA-1 is critical for its antiviral function. Mutant SVA-1 protein and splice variants lacking a nuclear export signal in the C-terminal domain show loss of antiviral activity. Full length SVA-1 protein contains both NLS and NES sequences suggesting it can shuttle back and forth from cytoplasm to nucleus. This process could be regulated by nuclear export proteins CRM1 (Exportin 1) and RAN^28,29^. HIV-1 protein Rev utilizes these nuclear export proteins for transferring unspliced and singly spliced HIV-1 transcripts to the cytoplasm from the nucleus. It is an interesting hypothesis to investigate the competitive inhibitory function of SVA-1 at this crucial stage of HIV-1 life cycle.

The precise mechanism of SVA-1 mediated inhibition of HIV-1 replication is requires further investigation. It will be interesting to study the SVA-1 effect on HIV regulatory proteins Tat and Rev in the context of viral genome transcription as well as export of un-spliced and singly-spliced viral RNA to the cytoplasm. Furthermore, SVA-1 protein may execute the function with direct or indirect interaction with viral and cellular proteins. Identification of interacting protein partners will shed light of the antiviral mechanism of the interferon stimulated SVA-1 factor. In summary, we have discovered the novel effector function of C19ORF66 as an inhibitor of HIV-1 replication.

## ACKNOWLEDGMENTS

This work was supported by a Cedars-Sinai Medical Center Programmatic Research Award to V.A., as well as a grant from the AIDS Healthcare Foundation to O.O.Y. We also thank Jason Nefalar for editing the manuscript.

## Author Contributions

V.A. and W.X. conceived and designed the experiments. W.X., J.I., D.C. and A.A. performed the experiments in various aspects of the project; V.A. O.Y., A.A., and W.X. analyzed the data. W.X., O.Y., and V.A. wrote the paper.

## EXTENDED DATA TABLES

**Extended Data Table 1.**
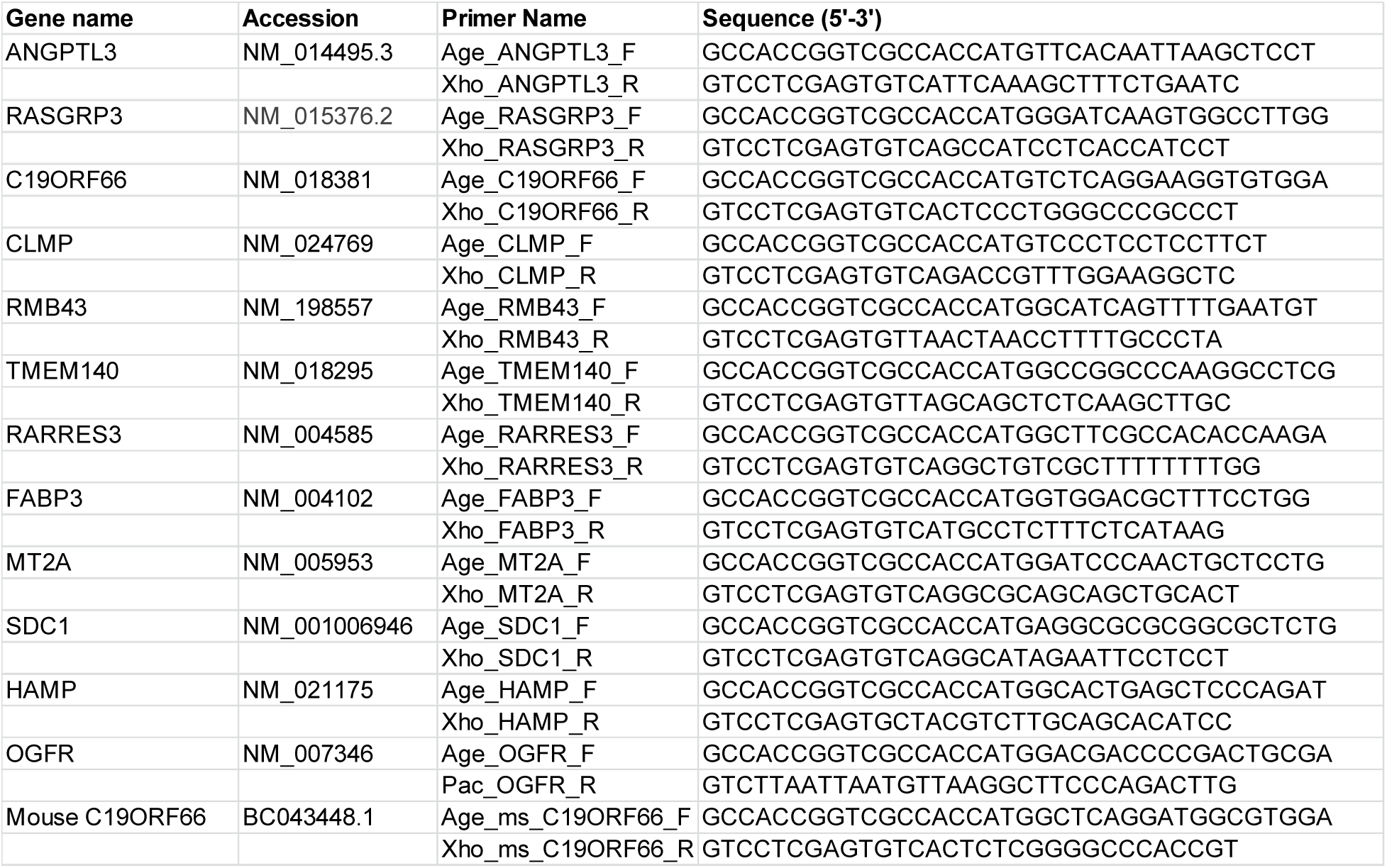
Primers used in this study.

**Extended Data Table 2.**
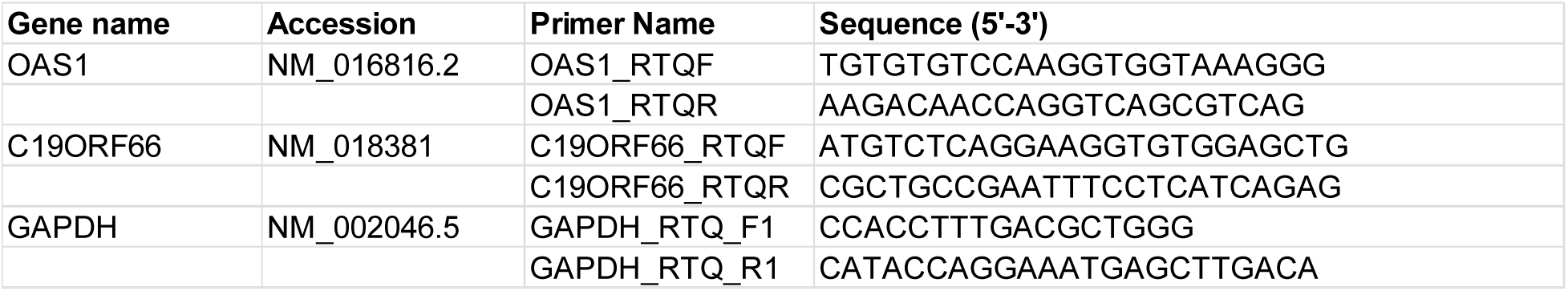
QPCR primers used in this study

## EXTENDED DATA FIGURES

**Extended Data Fig. 1.**
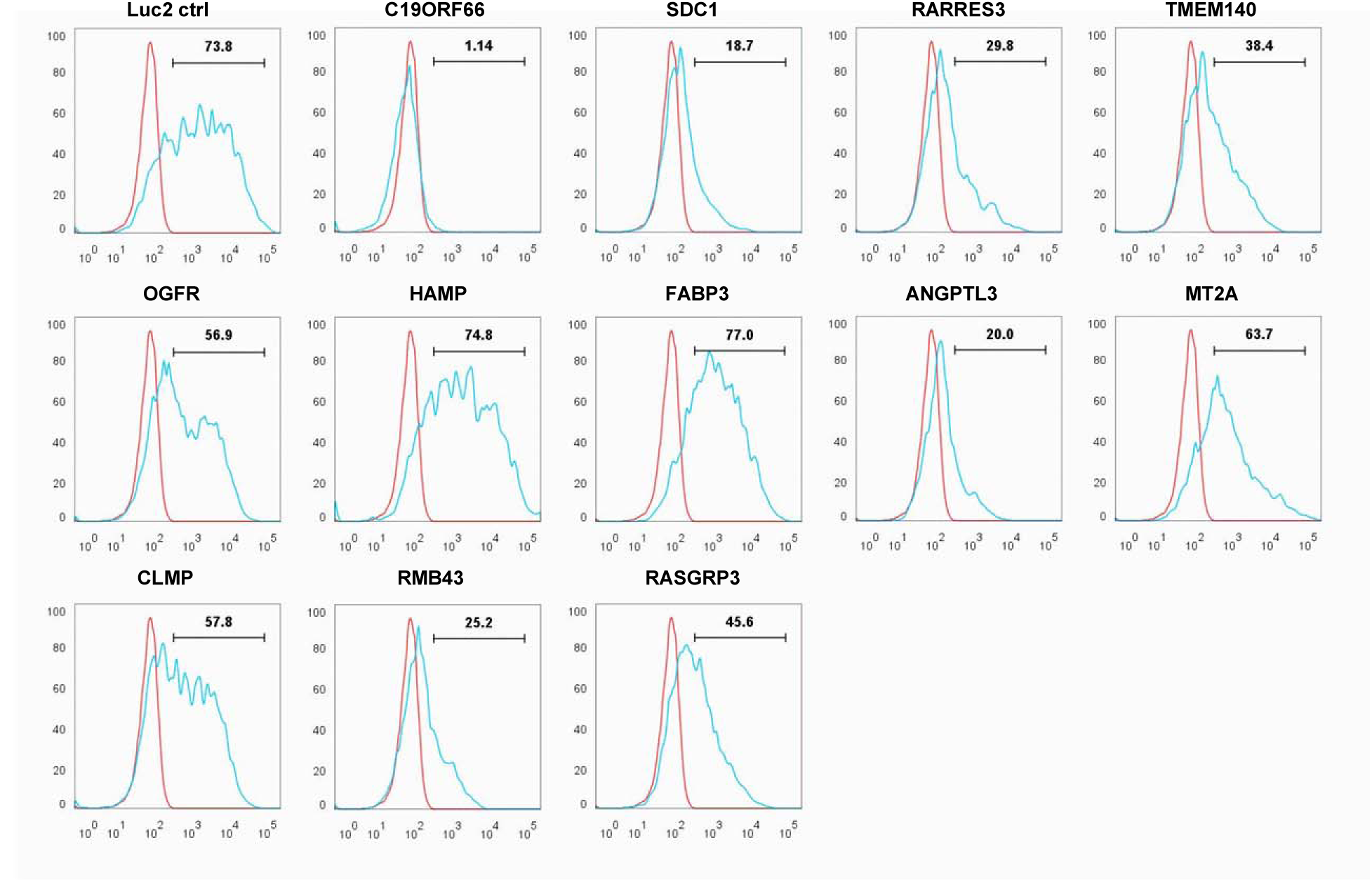
Flow cytometry based functional analysis of lentiviral particles. 293-FT cells transduced with indicated gene carrying lentivector were analyzed for number of mCherry fluorescent protein expressing cells at 48 hours post-transduction.

**Extended Data Fig. 2.**
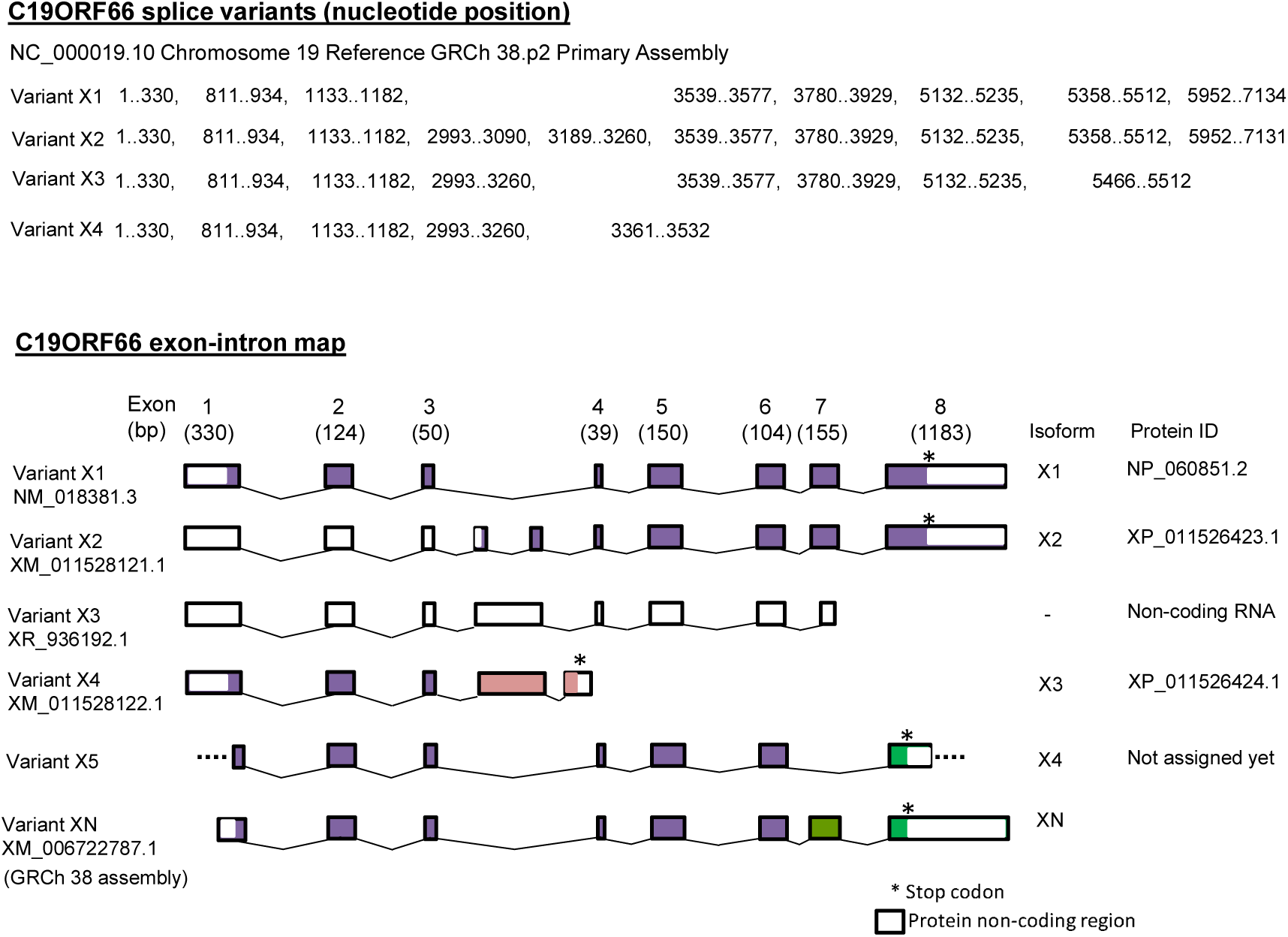
Splice variant analysis

C19ORF66 Isoform X4 sequence

MSQEGVELEKSVRRLREKFHGKVSSKKAGALMRKFGSDHTGVGRSIVYGVKQKDGQELSNDLDAQDPPEDMKQDRDIQAVATSLLPLTEANLRMFQRAQDDLIPAVDRQFACSSCDHVWWRRVPQRKEVSRCRKCRKRYEPVPADKMWGLAEFHCPKCRHNFRAPRAWDILCSPQEPEAEPPAQSAPPQQ PSH

C19ORF66 Isoform XN sequence

MSQEGVELEKSVRRLREKFHGKVSSKKAGALMRKFGSDHTGVGRSIVYGVKQKDGQELSNDLDAQDPPEDMKQDRDIQAVATSLLPLTEANLRMFQRAQDDLIPAVDRQFACSSCDHVWWRRVPQRKEVSRCRKCRKRYEPVPADKMWGLAEFHCPKCRHNFRPARPQGLGTDGVPVPLLRVRLPRVSNTDPPPALGPGPGSPQHPHSLLLSCRLLQPARAPRAWDILCSPQEPEAEPPAQSAPPQQPSH

**Extended Data Fig. 3.**
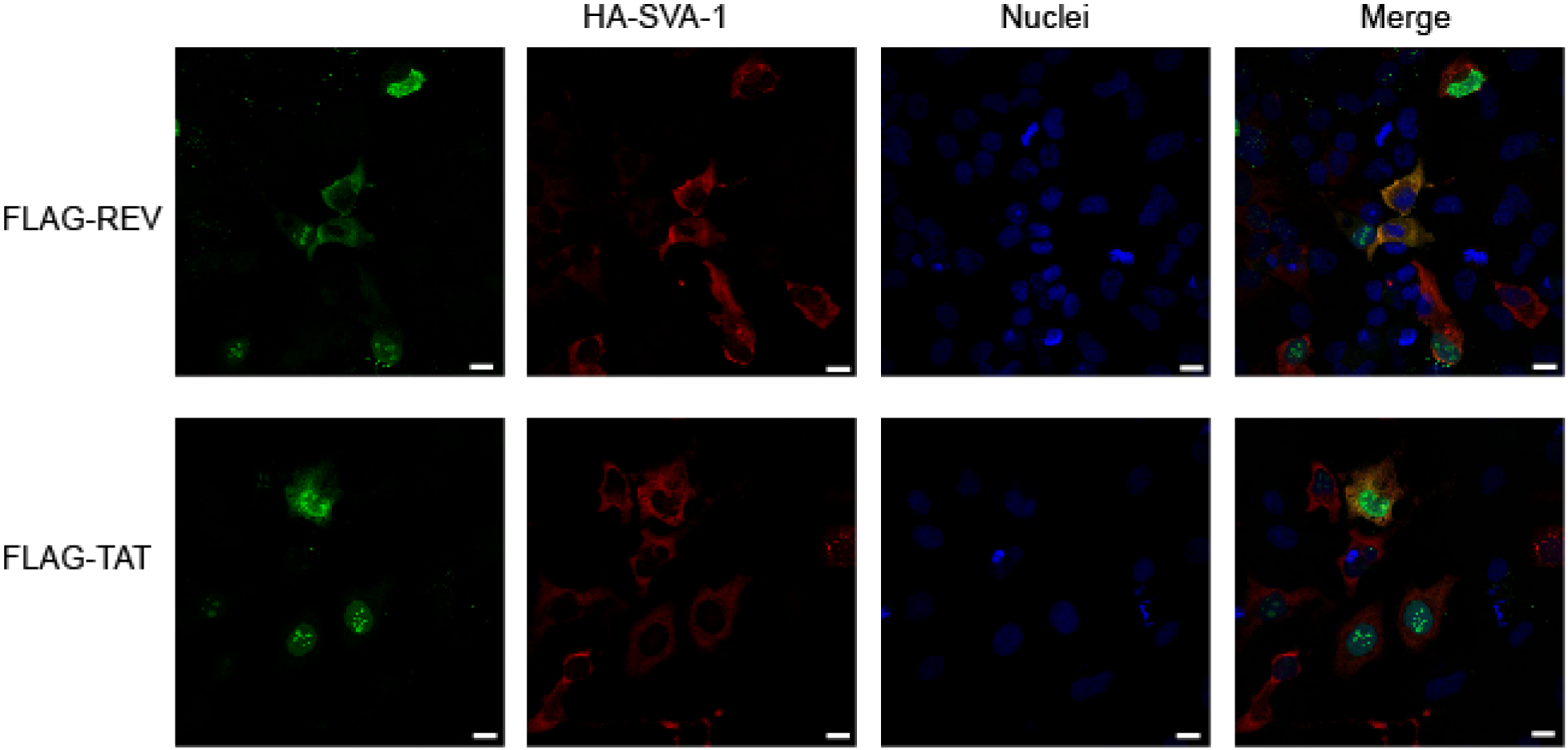
Confocal microscopic analysis of co-localization pattern. Full length SVA-1 protein shows occasional cytoplasmic co-localization with HIV-1 proteins REV and TAT. Both REV and TAT were predominantly found in the nucleus. Scale bar: 20 μm.

**Extended Data Fig. 4.**
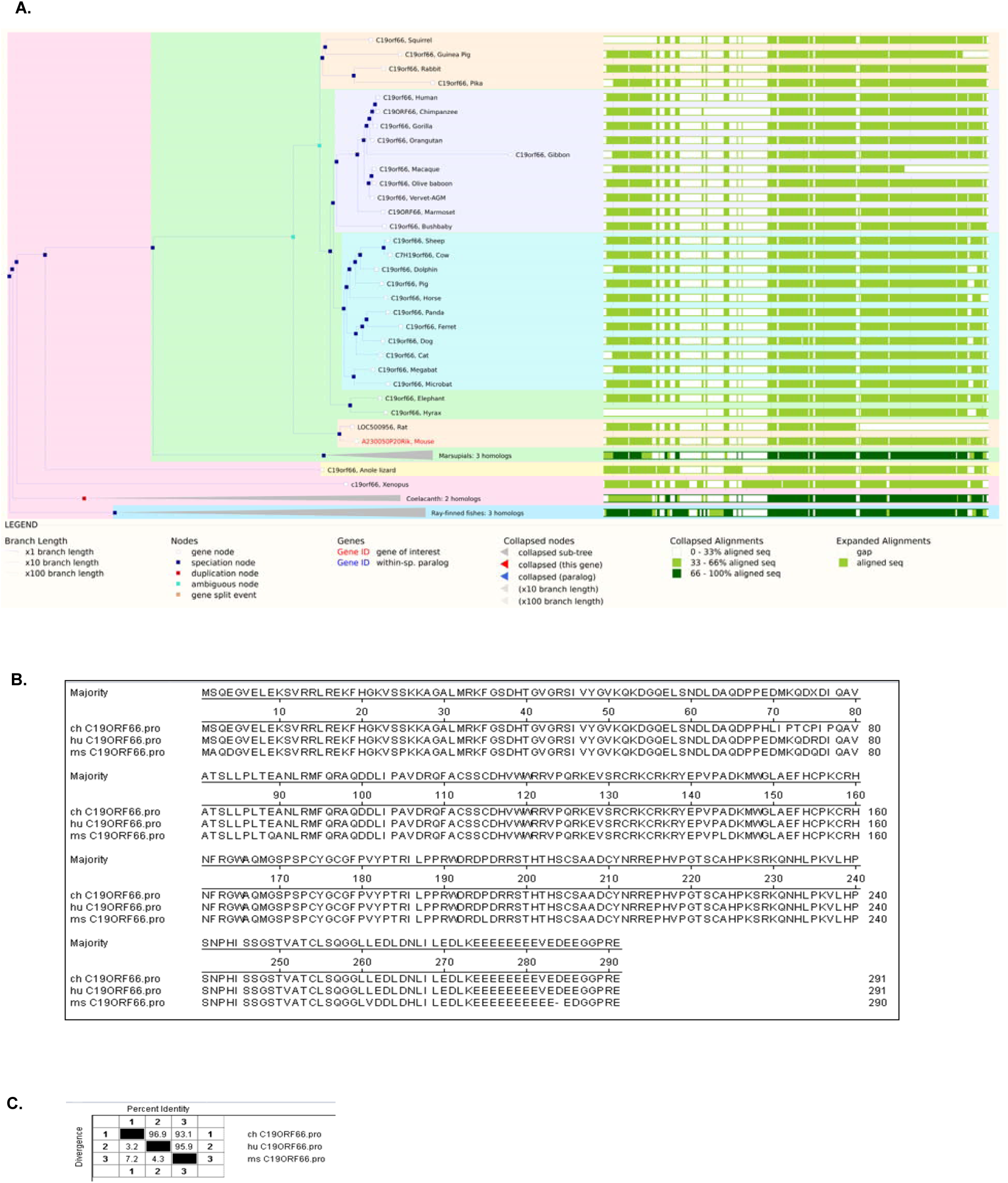
C19ORF66 is a conserved protein. (A) EnsEMBL alignment report of C19ORF66 protein sequence from different species (EnsEMBL_Web_Component_Gene_ComparaTree-Mus_musculus-Gene-Compara_Tree-79-ENSMUSG00000038884-10). (B) Phylogenetic tree of C19ORF66 and the Sequence Distances of C19ORF66 Alignment. (C) Table showing percent identity and divergence among the C19ORF66 proteins of various species

